# S100A8 and S100A9, biomarkers of SARS-Cov2-infected patients, suppress HIV replication in primary macrophages

**DOI:** 10.1101/2021.10.20.464686

**Authors:** Raphael M. Oguariri, Terrence W. Brann, Joseph W. Adelsberger, Qian Chen, Suranjana Goswami, Anthony R. Mele, Tomozumi Imamichi

## Abstract

S100A8 and S100A9 are members of the Alarmin family; these proteins are abundantly expressed in neutrophils and form a heterodimer complex. Recently, both proteins were identified as novel biomarkers of SARS-CoV-2 infection and were shown to play key roles in inducing an aggressive inflammatory response by mediating the release of large amounts of pro-inflammatory cytokines, called the “cytokine storm.” Although co-infection with SARS-CoV-2 in people living with HIV-1 may result in an immunocompromised status, the role of the S100A8/A9 complex in HIV-1 replication in primary T cells and macrophages is still unclear. Here, we evaluated the roles of the proteins in HIV replication to elucidate their functions. We found that the complex had no impact on virus replication in both cell types; however, the subunits of S100A8 and S100A9 inhibits HIV in macrophages. These findings provide important insights into the regulation of HIV viral loads in SARS-CoV2 co-infection.

## Introduction

S100A8 and S100A9 also known as myeloid-related protein (MRP) 8 and MRP14 are members of the S100 calcium binding protein family and belong to alarmin family protein [1,2]. They are low-molecular-weight (9–14 kDa) acidic proteins with important regulatory functions, including regulation of calcium buffering, cell differentiation, proliferation, cytoskeletal–membrane interactions, embryogenesis, cell migration, and inflammation [3-5]. The expression of human S100A8 and S100A9 are associated with acute and chronic inflammatory diseases [5-7]. S100A8 and S100A9 are expressed as homodimers or heterodimer complexes (S100A8/A9) and are constitutively expressed in neutrophils and monocytes [1,2,7,8]. They are not detected in resting tissue macrophages and lymphocytes but induced in macrophages upon stimulation [7,9]. S100A8, S100A9 and S100A8/A9 are chemoattractants for neutrophils [10] and stimulate the neutrophils’ adhesion to fibronectin [11]. It is also reported that S100A9 and S100A8/A9 enhance monocyte transmigration across endothelial cells [12].

During inflammation, the concentration of S100A8 and S100A9 in serum may rise up to near 100 μg /mL at local sites of inflammation [13]. Moreover, elevated serum levels of S100A8/A9 have recently been reported in patients infected with severe acute respiratory syndrome coronavirus 2 (SARS-CoV-2). Additionally, the S100A8/A9 complex released from neutrophils is considered a biomarker of Covid-19 infection [14,15] and is involved in the induction of cytokine storms [16]. It is reported that S100A8 and S100A9 complex is an endogenous ligand of Toll-like receptor 4 (TLR4) on dendritic cells (DC) and a recent study using primary dendritic cells demonstrated that cytosolic S100A9 suppresses HIV replication by inhibiting reverse transcription [17]. On contrary, S100A8 or S100A9 have been reported as HIV inducers/activators in studies using T cell lines or promonocytic cell line [18,19], however, a role of each S100A protein on HIV replication in primary cells were not investigated well yet.

Co-infection with SARS-CoV2 in people living with HIV (PLWH) with poorly controlled viral load may result in an immunocompromised status. Although the increase of S100A8/A9 in plasma is reported, the impact of extracellular S100A8/A9 on HIV replication in primary cells has not been thoroughly investigated. Therefore, in this study, we assessed the roles of S100A8 and S100A9 in HIV replication in primary macrophages and T cells.

## Materials and Methods

### Cells and Reagents

Peripheral blood mononuclear cells (PBMCs) were isolated from peripheral blood of healthy donors as previously described [20], after we obtained approval from the National Institute of Allergy and Infectious Diseases Institutional Review Board. CD4^+^ T cells and CD14^+^ monocytes were isolated from PBMCs using CD4 and CD14 beads (Miltenyi Biotech, Auburn, CA, USA), respectively. PBMCs and CD4^+^ T cells were stimulated with 5 μg/ml of PHA (Sigma Aldrich, St Louis, MO, USA), and CD14^+^ monocytes were differentiated into macrophages (MDMs) using Macrophage-colony stimulatory factor (M-CSF) (R&D systems, Minneapolis, MN, USA) as previously described [20]. Recombinant human S100A8 and S100A9 were obtained from Sino Biological Inc. (Beijing, China) and S100A8/S100A9 complex was purchased from R&D systems. Anti-SPTBN and anti-b-actin antibodies were obtained from Abcam (Waltham, MA, USA) and Sigma Aldrich, respectively.

### HIV replication assay

The HIV stocks of X4 (HIV_NL4.3_) [21] and R5 (HIV_AD8_) [22] viruses were prepared by transfection of the plasmid encoding each HIV strain into HEK293T cells, and the viruses were then purified from culture supernatants of the transfected cells using ultracentrifugation with 20% sucrose at 100,000 × _g_ for 2h and infectious titers were determined by the endpoint assay as previously described [23].The HIV replication assay was performed as previously described [20,24]. Briefly, PHA-stimulated CD4^+^ T cells were infected with X4 HIV-1 (MOI=0.001), and MDMs were infected with R5 HIV-1 (MOI=0.05). The infected PBMCs and CD4+ T cells were cultured at 37ºC in humidified air with 5% CO_2_ in the presence of 20 U/mL of IL-2 (Roche Diagnosis, Indianapolis, IN, USA), with various concentrations of S100A8 or S100A9 in RPMI-140 (Thermo Fisher, Waltham, MA) containing 10% fetal bovine serum (FBS; Cytiva, Marlborough, MA) [23]. HIV-infected MDMs were cultured for 14 days in Dulbecco’s modified Eagle medium (Thermo Fisher) containing 10% FBS. Culture medium was changed every three to 4 days with 50% concentration of S100 protein. The level of HIV-1 replication was determined by measuring p24 antigen in the culture supernatant using a kit (Perkin Elmer Inc, Shelton, CT, USA).

### Quantitation of HIV-1 Proviral DNA Copy Number

MDMs were treated or untreated with 1 μg/mL of S100A8 or S100A9 and cultured for 24 h at 37 °C. Pretreated cells were infected with DNase I-treated HIV_Ad8_ virus for 2 h at 37 °C. Cells were washed and cultured for additional 24 h. Absolute numbers of copies of proviral DNA were determined by real time PCR using custom made probes (Thermo Fisher). To obtain absolute proviral DNA copy numbers, standard curves were generated using serial dilutions of a plasmid encoding HIV gag region (pHIVgag) and RNaseP gene (pRNaseP) as previously described [24,25].

### FACS analysis

MDMs were cultured at 37 °C in humidified air with 5% CO_2_ for 24 h in the presence of media alone, 1 μg/ml of S100A8 or S100A9. After pretreatment, cells were washed with cold PBS, and then stained with fluorescein isothiocyanate (FITC)-conjugated anti-CD4 (BD-PharMingen, San Diego, CA, USA), or PE conjugated anti-CCR5 or anti-CD4 as previously described [26]. The stained cells were analyzed by using a Coulter XL flow cytometer (Beckman-Coulter, Fullerton, CA, USA). Events were collected by gating on forward and 90° light scatter. Positive fluorescence was determined using a matched isotype control IgG conjugated with FITC or PE (BD-PharMingen).

### HIV binding Assay

MDMs cells were pretreated with or without 1 μg/ml of S100A8 or S100A9 for 24 h. Cells were then incubated with DNase-treated HIV-1 at 4 °C for 2 h, followed by washing with ice-cold PBS. Total RNA was isolated using RNeasy Kit (Qiagen, Germantown, MD, USA) and reverse transcribed as previously described [24]. HIV-1 binding amount was evaluated by measuring HIV gag gene expression by real time PCR as described previously [24].

### Western blotting

Western blotting was performed as previously described [27]. MDMs were treated with or without 1 μg/ml of S100A8, S100A9 or 10 ng/mL of LPS for 2 days. Cells were lysed in radioimmunoprecipitation assay (RIPA) buffer(Boston, MA USA) supplemented with protease inhibitors (Sigma-Aldrich) and phosphatase inhibitor (Thermo Scientific, Rockford, IL). Protein amounts were determined using the BCA protein assay kit (Thermo Scientific). A total of 20 μg of whole cell lysate were resolved on a 3-8% Tris-Acetate NuPAGE (Invitrogen) and analyzed by immunoblotting with anti-SPTBN1, anti-APOBEC3G, (Abcam, Cambridge, MA, USA), or anti-Beta-Actin (Santa Cruz) as previously described [27]. Signals on the membranes were detected using HRP-conjugated anti-rabbit or anti-mouse immunoglobulins (GE Healthcare) with an ECL-plus kit (GE Healthcare). The intensity of the bands was analyzed by NIH Image J (http://rsbweb.nih.gov/ij/).

### Statistical analysis

Statistical analysis was performed using the Analysis ToolPak from Microsoft Excel. Results were representative of at least three independent experiments. The values were expressed as mean and SD of individual samples. Statistical significance was determined by the Student’s t test. A p-value < 0.05 was considered a significant difference between the experimental groups.

## Results and Discussion

To assess the function of each protein in HIV replication in primary cells, the activated CD4^+^-T cells and MDMs were infected with X4 HIV-1 and R5 HIV-1, respectively [23]. The infected cells were cultured at 37°C under physiological concentrations (0–10 μg/mL) of recombinant S100A8/A9 by exchanging half of the medium every 3–4 days with fresh medium containing the same concentration of protein[23]. As shown in Fig. 1a and 1b, the S100A8/A9 complex had no impact on HIV replication in CD4^+^ T cells and MDMs. S100A8 and S100A9 have been reported to induce HIV in T cell lines [19]; thus, we assessed the impact of each recombinant protein on HIV replication in primary T cells and MDMs. HIV-infected T cells and MDMs were cultured in the presence of different concentration of S100A8 or S100A9. Interestingly, although both proteins had no effect on HIV replication in primary T cells (Fig. 1c), they inhibited HIV replication in MDMs in a dose-dependent manner (Fig. 1d). In the presence of 10 μg/mL S100A8 and S100A9, HIV replication in MDMs was suppressed by 89±10% and 79±5.4% (n=3, p<0.05), respectively, indicating that S100A8 and S100A9 exhibited anti-HIV effects in primary MDMs.

**Figure 1.**
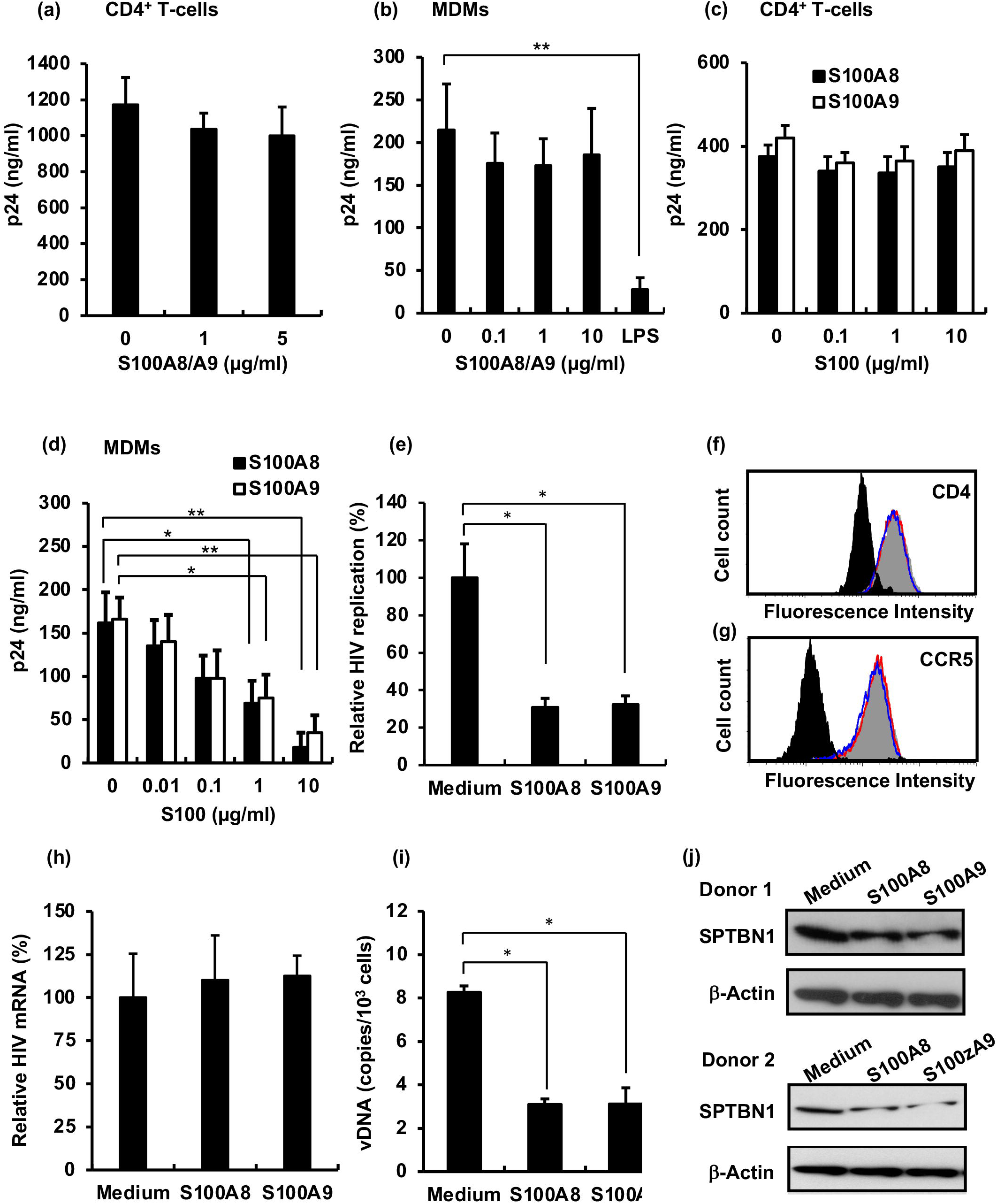
Evaluation of anti-HIV effect of S100A8/A9, S100A8 and S100A9 in primary cells. (a-b) T-cells (a) and MDMs (b) were infected with HIV-1 and then cultured in the presence of different concentrations of S100A8/A9 combination. As a positive control of TLR4 stimulation, 10 ng/ml of LPS (Sigma Aldrich) was used. Each assay was performed in quadruplicate and half of the culture supernatants were changed in all experiments on every 3^rd^ or 4^th^ day with the fresh media containing the same concentration of the recombinant proteins. HIV-1 replication was monitored HIV p24 antigen in the culture supernatant by a p24 antigen capture assay. Data show means ± SD and are representatives of three independent experiments. (c and d) HIV-infected T cells (c) and MDMs (d) were cultured in the presence of different concentrations of S100A8 or S100A9. Each assay was performed in quadruplicate. HIV-1 replication was monitored HIV p24 antigen in the culture supernatant by a p24 antigen capture assay. Data show means ± SD. and are representatives of three independent experiments. (e) MDMs were pulse-treated with S100A8 or S100A9 for 24 hr, and then the treated cells were infected with HIV. The infected MDMs were culture for 14 days in the absence of S100 proteins with changing fresh medium on every 3^rd^ or 4^th^ day. HIV-1 replication was monitored HIV p24 antigen in the culture supernatant by a p24 antigen capture assay. Data show means ± SE (n=3). (f and g) CD4 (f) and CCR5 (g) expression on the treated cells were compared to that on untreated cells. Back shows isotype control, gray, red and blue histogram indicate untreated cells, S100A8 and S100A9-treated cells, respectively. Data shows a representative of three independent experiments. (h) The binding assay was performed using the pretreated MDMs. The cells were incubated with HIV on ice for two hr to allow virus binding. Virus RNA on cell surface was determined by qRT-PCR. The data show the means ± SE of three independent experiments. (i) MDM cells were pretreated with S100A8 and S100A9 for 24 h at 37°C. Cells were washed and infected with HIV for 2 h. The cells were washed and then cultured for an additional 24 h in the absence of recombinant proteins. The genomic DNA was isolated and used in qPCR for quantification of proviral DNA copy numbers. A standard curve was generated using serial dilutions of plasmids encoding HIVgag or RNaseP gene to obtain absolute proviral DNA copy numbers as previously described[23]. The results represent DNA copy numbers per 1 × 10^6^ cells. The data show the means ± SE from three independent experiments. (j and k) Expression levels of SPTBN1 in MDMs in whole-cell lysates from two independent donors’ cells were compared by Western blotting using an anti-SPTBN1 antibody (Abcam)[23]. As an internal control, anti-b-actin (Sigma Aldrich) was used. The intensity of Western blot bands was analyzed using the ImageJ (NIH). Statistical analysis was done using Prism (GraphPad). Statistical significance was determined by the Student’s t test. A p value<0.05 was considered a significant difference. * p<0.05, **p<0.01. Short version

To further characterize these anti-HIV effect, we pulse-treated MDMs with 1 μg/mL of each protein for 24 h and then infected the pre-treated cells with HIV. Infected cells were cultured in the absence of S100 proteins and viral replication was monitored. Monitoring of viral replication (Fig. 1e) showed that the S100A8- and S100A9-treatment were sufficient to suppress HIV replication by 65±9.9% and 77±6.7% (p<0.05, n=3), respectively. This inhibitory effect may involve suppression of HIV infection or interference with the viral cycle after infection. To further elucidate the inhibitory mechanism, we first compared the expression levels of CD4 and CCR5 in pulse-treated cells using FACS, as previously described. FACS analysis demonstrated no changes in CD4 (Fig. 1f) or CCR5 expression (Fig. 1g). Thus, we assumed that the inhibition was caused by suppression of virus binding to receptors. Consequently, we conducted HIV binding assays using qRT-PCR. Pretreated cells were incubated with HIV for 2h at 4°C. Total RNA (cellular RNA and viral RNA on the cell surface) was extracted, and qRT-PCR was then conducted. HIV gag viral RNA levels were normalized to *GAPDH* mRNA levels, and the amount of HIV RNA was then compared. Pre-treatment had no effect on HIV binding (Fig. 1h). Next, we assessed the inhibitory effects of reverse transcription by measuring the copy numbers of proviral DNA using qPCR. HIV-infected cells were cultured for 24 h, and genomic DNA was extracted and subjected to qPCR as previously described [23]. S100A8 and S100A9-treatment decreased proviral DNA copy numbers by 63± 3.1% and 63±9.2 % (p<0.01, n=3) (Fig. 1i), suggesting that pre-treatment suppressed HIV replication during reverse transcription after infection. We previously reported that Spectrin β Non-Erythrocytic 1 (SPTBN1) plays a key role on HIV replication in MDMs and downregulation of the expression of SPTBN1 suppressed HIV reverse transcription [23]. To address a potential mechanism of S100A8 or S100A9-mediated RT suppression, we conducted Western blotting using cell lysates from the pre-treated cells and anti-SPTBN1 antibody [23]. S100A8- and S100A9-treatment demonstrated a partial downregulation of SPTBN1 expression to 58±24% and 60±28% (p>0.05, n=3), respectively (Fig. 1j), indicating that SPTBN1 may be involved in HIV inhibition in the pretreated cells. HIV reverse transcription is regulated by many host-required factors [23]; thus, further studies are needed to elucidate the mechanisms mediating these inhibitory effects.

The concentrations of S100A8/A9 proteins are increased in patients with SARS-CoV2 infection [14], and these proteins are involved in the cytokine storm. Thus, we speculated that elevation of the concentrations of both proteins may affect HIV replication in patients with HIV infection in whom the disease is poorly controlled by combination antiretroviral therapy because HIV replication was shown to be enhanced by S100A8 and S100A9 in a previous report [19]. However, our data indicated that the complex proteins did not alter HIV replication in primary cells, suggesting that although the protein concentration was increased in PLWH co-infected with SARS-CoV2, these proteins may not directly enhance HIV replication and instead each subunit may inhibit HIV replication. S100A9 is an intracellular HIV inhibitor that suppresses the HIV reverse transcription reaction[17], and our data demonstrated that the protein is an extracellular inhibitor; therefore, S100A9 may have applications as therapeutic reagents. Impact of S100A8 and S100A9 in co-infection with SARS-CoV2 and other viruses, such as influenza virus or Hepatitis C virus, should also be considered, and further studies may provide important insights into the regulation of viral load in SARS-CoV2 co-infection.

## Acknowledgments

All authors thank Dr. HC. Lane for supporting this project and Drs. W. Cheng, H. Sui and S. Laverdure for critical reading. The plasmid encoding HIV_NL4.3_ and HIV_AD8_ gene were obtained from Dr. M. Martin (NIAID, Bethesda, MD, USA).

## Authorship confirmation

All authors read the final version of the manuscript and agreed to the context in the manuscript.

## Conflicts of Interest

All authors declare that there are no conflicts of interest.

## Funding Status

This project has been funded in whole or in part with federal funds from the National Cancer Institute, National Institutes of Health, under Contract No. HHSN261200800001E. The content of this publication does not necessarily reflect the views or policies of the Department of Health and Human Services, nor does mention of trade names, commercial products, or organizations imply endorsement by the U.S. Government. This research was supported [in part] by the National Institute of Allergy and Infectious Disease. The content of this publication does not necessarily reflect the views or policies of the Department of Health and Human Services, nor does mention of trade names, commercial products, or organizations imply endorsement by the U.S. Government.

## Notes

### Competing Interest Statement

The authors have declared no competing interest.

